# Unearthing Regulatory Axes of Breast Cancer circRNAs Networks to Find Novel Targets and Fathom Pivotal Mechanisms

**DOI:** 10.1101/569756

**Authors:** Farzaneh Afzali, Mahdieh Salimi

**Affiliations:** Medical Biotechnology Department, National Institute of Genetic Engineering and Biotechnology (NIGEB), Tehran, Iran; Pars Silico Bioinformatics Laboratory, Tehran, Iran

**Keywords:** circRNA, miRNA, breast cancer, regulatory network, microarray analysis

## Abstract

Circular RNAs (circRNAs) along other complementary regulatory elements in ceRNAs networks possess valuable characteristics for both diagnosis and treatment of several human cancers including breast cancer (BC). In this study, we combined several systems biology tools and approaches to identify influential BC circRNAs, RNA binding proteins (RBPs), miRNAs, and related mRNAs to study and decipher the BC triggering biological processes and pathways.

Rooting from the identified total of 25 co-differentially expressed circRNAs (DECs) between triple negative (TN) and luminal A subtypes of BC from microarray analysis, five hub DECs (hsa_circ_0003227, hsa_circ_0001955, hsa_circ_0020080, hsa_circ_0001666, and hsa_circ_0065173) and top eleven RBPs (AGO1, AGO2, EIF4A3, FMRP, HuR (ELAVL1), IGF2BP1, IGF2BP2, IGF2BP3, EWSR1, FUS, and PTB) were explored to form the upper stream regulatory elements. All the hub circRNAs were regarded as super sponge having multiple miRNA response elements (MREs) for numerous miRNAs. Then four leading miRNAs (hsa-miR-149, hsa-miR-182, hsa-miR-383, and hsa-miR-873) accountable for BC progression were also introduced from merging several ceRNAs networks. The predicted 7- and 8-mer MREs matches between hub circRNAs and leading miRNAs ensured their enduring regulatory capability. The mined downstream mRNAs of the circRNAs-miRNAs network then were presented to STRING database to form the PPI network and deciphering the issue from another point of view. The BC interconnected enriched pathways and processes guarantee the merits of the ceRNAs networks’ members as targetable therapeutic elements.

This study suggested extensive panels of novel covering therapeutic targets that are in charge of BC progression in every aspect, hence their impressive role cannot be excluded and needs deeper empirical laboratory designs.

## INTRODUCTION

Breast cancer (BC) is the most lethal cancer among women worldwide in 2017 [1]. To surpass the anti-cancer treatments’ deficiencies, exploitation of all the novel high throughput data rooting from underlying microlayers of a disease is critical. Competing endogenous RNAs (ceRNAs) network which consists of various transcripts regulating each other at post-transcriptional level [2], is a complex and yet novel tracing means to investigate the breast cancer progression and metastasis. The interrelation between the transcripts that possess miRNA response elements (MREs), namely circular RNAs (circRNAs) and mRNAs is pivotal in the deciphering steps. The fundamental core of the ceRNA network is based on the ratio of affinity and competing for shared miRNAs MREs [3].

The circRNAs that were surmised to be non-coding, are now believed to be translated into proteins [4]. Apropos of being resistant to exonucleases and being more stable due to the lack of cap and poly A tales in their structure [5], possess myriad noteworthy functions in genes’ expression regulations in cancer. They are divided into three subcategories regarding their source of emergence; exonic circRNA (ecRNA), exon-intron circRNA (ElciRNA), and circular intronic circRNA (ciRNA) [6-8]. In parallel with functioning as transcription and translation regulator, the ecRNAs act as miRNAs sponges enjoying MREs [9]. The circRNA profile of BC versus normal tissues introduces manifold anti-breast cancer controllable targets [10]. Suppression of circIRAK3 leads to down-regulation of *FOXC1* through miR-3607-FOXC1 axes and promotes BC cell metastasis [11]. The proliferation and apoptosis of BC cells would be ceased and enhanced respectively, following ABCB10-circRNA elimination [12]. These and other analogous studies give prominence to the emerging role of circRNAs in controlling BC development.

The miRNAs as the indispensable player of the ceRNAs network with a length of 22 nucleotides, attach to 3’ end of mRNAs and act as inhibitors [13]. OncomiRs, including tumor suppressors and tumor triggers, are one of the main targets of breast cancer therapies these days [14]. The overexpression of miR-21 and miR-155, and lost expression of miR-200 and let-7 are responsible for BC progression [15]. The microRNA-34a was authenticated to be a migration inhibitor by targeting EMT transcription factors in BC [16]. The cluster of mir-183/-96/-182 which is under the direct regulation of HSF2 and ZEB1 significantly overexpresses in BC subtypes as oncomiRs. Arresting this cluster triggers BC cell death and subsides BC cells migration and proliferation [17].

The regulation orchestra would be more complete by another important player scilicet RNA binding proteins (RBPs) that can be sponged by circRNAs. Post-transcriptional modification relies on non-coding RNAs and RBPs forming a complex layer of regulation that its disarrangement results in various hard to cure diseases including cancer [18]. As an omnipotent regulatory element, its range of activities differs from pre-mRNA related regulations to RNA turn over and localization, hence has a fundamental impact upon balancing any cellular mechanisms counting growth, cell death, immune recognition, and metastasis [19-21]. It was noted that RNA binding motif protein 38 (RBM38) as a tumor suppressor RBP, exerts it impact through regulating STARD13 ceRNA network and in return inhibits BC metastasis [22]. On the other hand, HuR that counts as tumor trigger enhances Cyclin-E expression and promotes BC progression [23].

In the current study, we took advantage of a coalescing investigation of both systems biology and microarray analysis in order to find differentially expressed circRNAs (DECs) in BC. To narrow down the discovered circRNAs, their corresponding genes were investigated in BC regarding their expression status via RNA-seq data available in Pan-Cancer Analysis Platform of starBase. The circRNAs that were identically expressed with their corresponding genes opted for further analyses. Furthermore, probable sponged miRNAs were predicted by Circular RNA Interactome database and filtered by miRCancer to have the curated differentially expressed BC miRNAs. The “circRNAs-miRNAs” (ceRNAs) network was formed and visualized in Cytoscape 3.6.0. The network then was investigated to find the hubs of circRNAs and miRNAs by NetworkAnalyzer tool of Cytoscape. The RBP profiling and super sponge confirmation of hub circRNAs were performed by Circular RNA Interactome database. The STRING database then was used to study the interactions of RBPs and their functional annotation. The TargetScan algorithm of DIANA-miRPath3 was then used to spot downstream mRNAs of miRNAs. The predicted mRNAs were filtered by differentially expressed genes (DEGs) identified through microarray analysis. The protein-protein interaction (PPI) network of remaining mRNAs was established via STRING database and the hub clusters were found by MCODE plugin of Cytoscape to form the “miRNAs-mRNAs” network. The hubs were then subjected to DAVID 6.7 to spot enriched GO terms and pathways. The BC leading miRNAs then were spotted through merging the formed networks and experienced other analyses including enrichment analysis and prediction of MREs that were employed by various promising computational biology tools.

## METHODS

### 1. Identifying BC DECs and DEGs Through Microarray Analysis

The expression dataset of BC circRNAs (GSE101123) was obtained from GEO (https://www.ncbi.nlm.nih.gov/geo/) database. The dataset contains two important subtypes of BC namely, triple negative (TN) and luminal A. The raw data was imported in RStudio 1.1.442 and analyzed by Limma package and applying RMA normalization algorithm. The |log (FC)| > 1 and *p-*value < 0.05 were the established criteria for finding DECs. The common co-expressed DECs between mentioned subtypes were selected for further studies.

The gene expression dataset of GSE65194 [24], which has been studied subtypes of TN and luminal A simultaneously, was selected to introduce the DEGs in BC. All the analysis steps were identical but the adjusted *p-*value less than 0.05 and also, the average expression was calculated in case of multiple probes genes. The common co-expressed DEGs were selected for further studies.

### 2. The Verification of Discovered DECs

Gene Differential Expression tool of starBase Pan-Cancer Analysis Platform (http://starbase.sysu.edu.cn/panCancer.php), having more than 10000 RNA-seq expression data across 32 types of cancer, was used to filter the DECs regarding their corresponding genes expression in BC. The DECs that were not co-expressed with their genes were removed.

### 3. The “circRNAs-miRNAs” Network Establishment and Hubs Identification

The DECs tied miRNAs were predicted by Circular RNA Interactome (https://circinteractome.nia.nih.gov/index.html) and then filtered by curated BC miRNAs of miRCancer database (http://mircancer.ecu.edu/). The “circRNAs-miRNAs” network then was constructed utilizing Cytoscape v3.6.0.

The “circRNAs-miRNAs” network was subjected to NetworkAnalyzer tool of Cytoscape and Degree, as the simplest centrality index, was calculated as the direct number of each gene’s neighbors to find the hub circRNAs and hub miRNAs. The hub circRNAs-miRNAs and hub miRNAs-circRNAs networks then were merged intersected together to form the “hub miRNAs-hub circRNAs” network.

### 4. RBP Profiling and Super Sponge Mapping

To identify the attachable RBPs of hub circRNAs, besides the ones which bind to circRNAs flanking regions and junctions, Circular RNA Interactome database was used [25]. Subsequently, the identified RBPs were subjected to STRING v10.5 (https://string-db.org/) database (interaction score: 0.4 and text mining was excluded) to form their interaction status and study their GO terms and related KEGG and Reactome pathways.

Then to find whether the discovered hub circRNAs also count as the super sponge in general or not, miRNAs target sites were explored through the same database.

### 5. Constructing PPI Network, Exploring Hub Nodes, and Enrichment Analysis

The miRNAs-mRNAs interactions were predicted through TargetScan algorithm of DIANA-miRPath 3 (http://snf-515788.vm.okeanos.grnet.gr/index.php?r=mirpath). Then the obtained mRNAs were filtered by discovered DEGs of the current study. The common ones were selected and subjected to STRING v10.5 database applying the 0.7 as the minimum required interaction score and excluding text mining as the active interaction source to form the PPI network. The resulted network then was imported into Cytoscape v3.6.0 and merged with BC luminal A and TN DEGs again to have their expression status as well.

The MCODE plugin of Cytoscape which finds the highly interconnected complexes of a network was used in order to investigate the hubs within clusters and form the “mRNAs-miRNAs” network. The clusters were then subjected to Enrichment Analysis (KEGG pathway and GO TERM analyses) tool of DAVID 6.7 database (https://david-d.ncifcrf.gov/).

### 6. Enrichment Analyses and MREs Exploration of leading miRNAs

The common (leading) miRNAs of “miRNAs-mRNAs” and “hub miRNAs-hub circRNAs” networks were sent to miRNA Differential Expression tool of Pan-Cancer Analysis Platform to monitor their expression status between normal and breast cancer tissues. The leading miRNAs were subjected to miRPathDB v1.1 (https://mpd.bioinf.uni-sb.de/overview.html) for Enrichment Analysis. The MREs prediction took place upon leading miRNAs and related hub circRNAs by Circular RNA Interactome database.

## RESULTS

### 1. Breast Cancer DECs and DEGs

The datasets information was listed in Supplementary 1(Sheet A). There were 79 and 175 total DECs for luminal A (50 up-regulated and 29 down-regulated circRNAs) and TN (90 up-regulated and 85 down-regulated), respectively. The lists were merged together to find the common co-expressed DECs. The resulted list contained 67 circRNAs including 41 up-regulated and 25 down-regulated ones. Following filtering the DECs’ list with RNA-seq data of circRNAs’ corresponding genes, the 25 DECs were identically expressed with their genes and selected for further analyses (Supplementary 1; sheets B & C).

The total DEGs of luminal A and TN subtypes were accordingly 2203 (1033 up-regulated and 1170 down-regulated) and 3148 (1622 up-regulated and 1526 down-regulated). After merging the lists, there were 1384 (553 up-regulated and 831 down-regulated) co-expressed DEGs left (Supplementary 1; sheet D).

### 2. Establishment of circRNAs-miRNAs network and Hubs Investigation

The total number of 247 miRNAs were predicted by Circular RNA Interactome database for the discovered DECs. After filtering the list with curated BC miRNAs of miRCancer database, the total number was decreased to 44 (Supplementary 2). The “circRNAs-miRNAs” network then was manifested in Cytoscape 3.6 harboring 70 nodes and 112 edges. The network was exposed to hubs investigation by NetworkAnalyzer tool. The 20 percent of each type nodes count as hubs. The total of 5 circRNAs (hsa_circ_0001666, hsa_circ_0001955, hsa_circ_0003227, hsa_circ_0020080, and hsa_circ_0065173) and 9 miRNAs (hsa-miR-145, hsa-miR-149, hsa-miR-182, hsa-miR-203, hsa-miR-31, hsa-miR-330-3p, hsa-miR-383, hsa-miR-520g, and hsa-miR-873) were derived to be the corresponding hubs. The networks then were merged intersected to form the “hub circRNAs-hub miRNAs” network with 14 nodes and 20 edges (Figure 1 and Table 1).

**Fig1.**
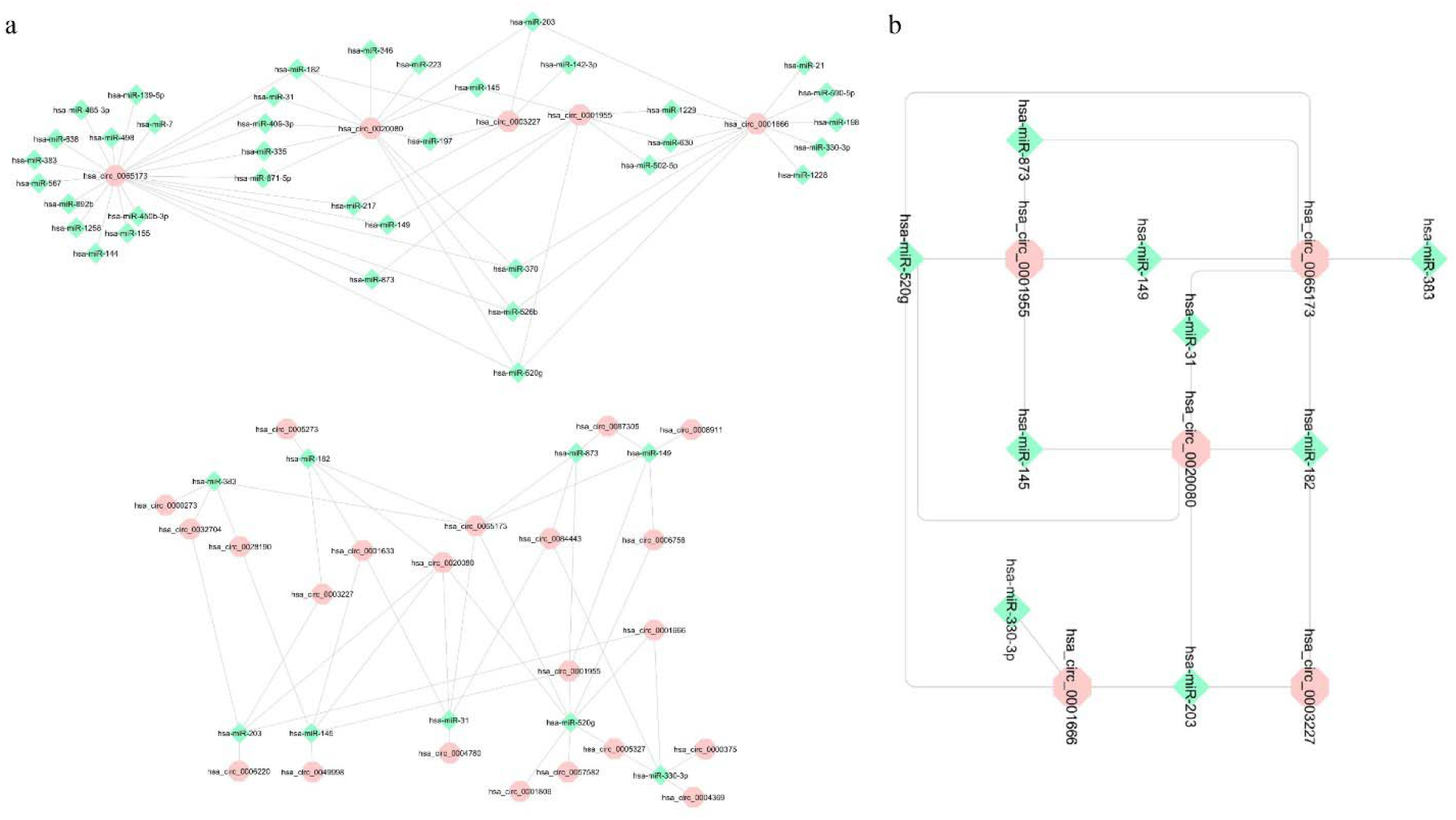
The identified hubs in circRNA-miRNA network. (a) The identified circRNA and miRNA hubs through NetworkAnalyzer tool of Cytoscape 3.6.0 were brought in separate networks. (b) The common elements between the found networks. The pink hexagon shaped and light green diamond shaped nodes are respectively hub circRNAs and hub miRNAs.

**Table 1.**
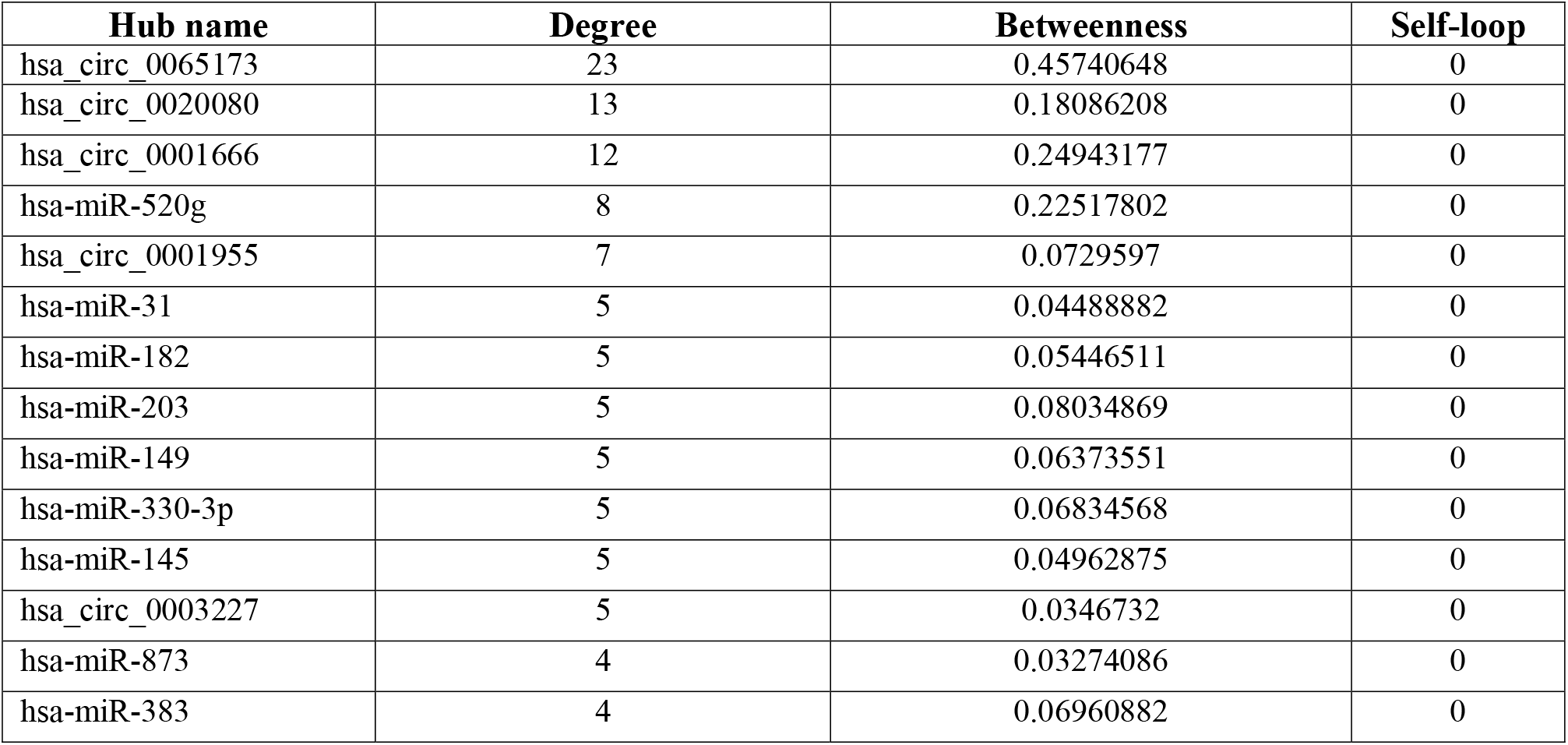
The found hub circRNAs and miRNAs. The hub elements (top 20% of all the elements) were excavated from circRNA-miRNA network via NetworkAnalyzer tool of Cytoscape 3.6.0 and ranked by their Degree scores indicating their neighbor nodes while not forming any self-loop.

### 3. RBP Profiling and Super Sponge Mapping

Following searching BRPs, a total of 26 ones were identified for the five hub circRNAs. The top five of each circRNA concerning binding sites were (11 ones in total) was plotted by ggplot2 package of RStudio (Figure 2(a)). Their interactions were also investigated by STRING and the related network contained 12 nodes (PTBP1 and PTBP2 were chosen for PTBP) and 17 edges (Figure 2(b)). The most enriched BP; biological process, CC; cellular component, MF; molecular function, and pathways (KEGG and Reactome) were respectively, negative regulation of translation [GO:0017148], ribonucleoprotein complex [GO:1990904], mRNA binding [GO:0003729], RNA transport and equally Transcriptional misregulation in cancer [hsa03013, hsa05202], and Insulin-like Growth Factor-2 mRNA Binding Proteins [HSA-428359] (Figure 2(c) and Supplementary 3(A)).

**Fig2.**
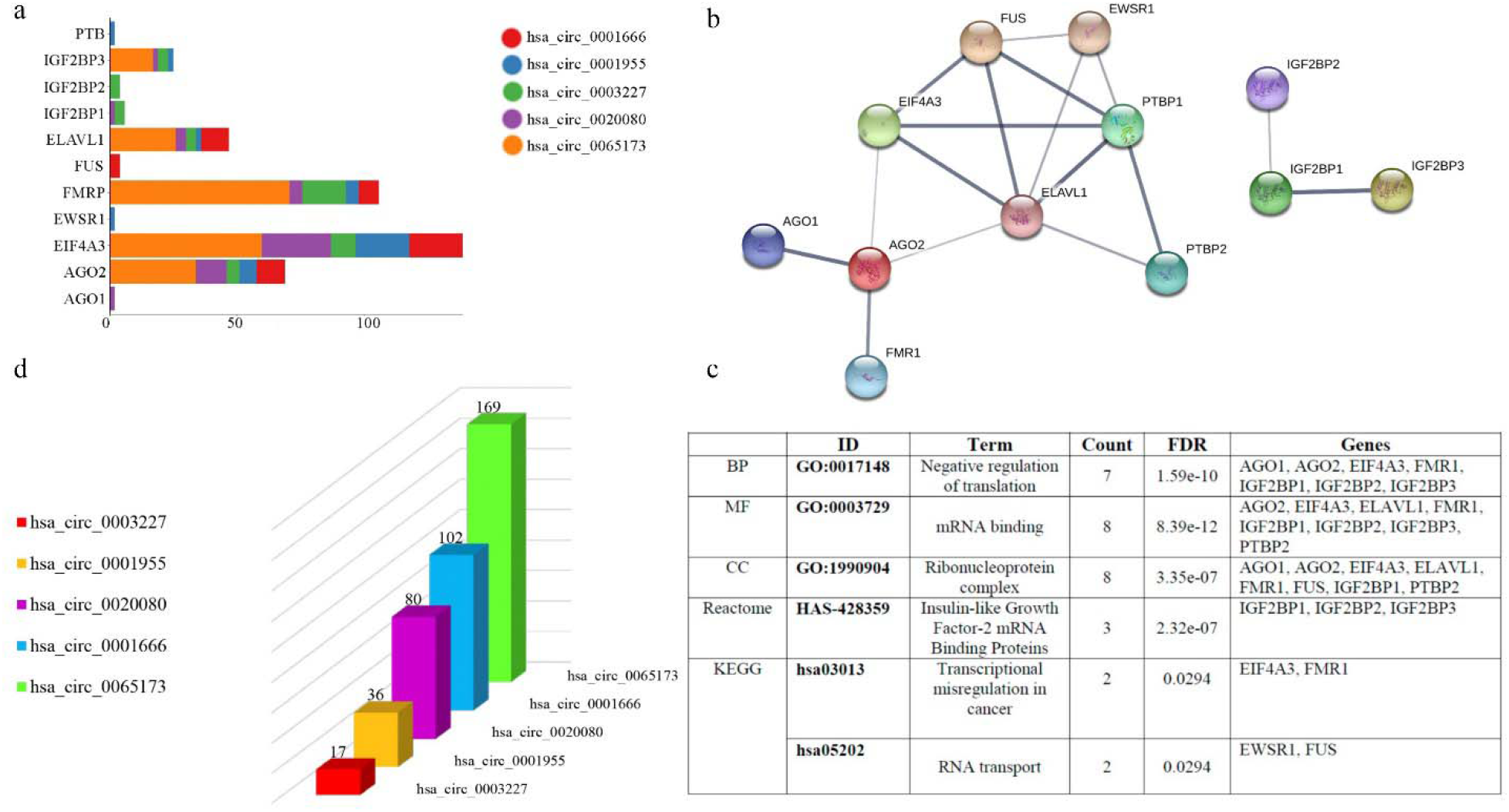
The RBPs profiling and mapping super sponge circRNAs. (A) The bar plot of hub circRNAs along their top 11 predicted RBPs (top 5 for each circRNAs). The Y-axis engrossed titles of RBPs and X-axis regarded as their count. Each color represents one type of circRNAs which were brought as legend of the plot. The plot was formed by ggplot2 package of R-studio. B) The excavated interactions of RBPs through STRING database. The thicker edges represent the higher probability of that interaction. The PTB were brought in names of PTBP1 and PTBP2. (C) The most enriched GO terms and pathways of top RBPs by STRING database along the false discovery rate (FDR) scores, count, and names of the responsible genes. (D) The identified miRNAs recognition sites on each hub circRNAs indicating that all of them count as super sponge circRNAs. Each color represent each circRNAs that were brought as legend on the left side of the graph that was made by Microsoft Excel 2013.

As the mapped miRNAs for hub circRNAs [hsa_circ_0003227, hsa_circ_0001955, hsa_circ_0020080, hsa_circ_0001666, and hsa_circ_0065173] were respectively, 17, 36, 80, 102, 169 ones, all of them count as super sponge possessing strong capability of gene expression regulation through engaging a lot of miRNAs (Figure 2(d) and Supplementary 3(B)).

### 4. Identified Hubs in the PPI regulatory network, Enriched GO Terms and Pathways

The DEGs filtration of TargetScan algorithm of DIANA-miRPath3 predicted miRNAs related mRNAs, resulted in 393 mRNAs (Supplementary 4; sheet A). With the purpose of exploiting their interactions profoundly, a PPI network (128 nodes and 178 edges) was established and visualized by Cytoscape 3.6.0 and after merging with DEGs the 119 nodes with 156 edges left. To narrow down the PPI network and due to the importance of hubs within clusters, the primary network divided to 7 clusters (Scores: 6, 5.556, 4, 3, 3, 3, and 3; overall: 32 nodes and 58 edges) utilizing MCODE plugin applying k-score=2. The resulted PPI network further combined with related miRNAs to form a precise “hub mRNAs-miRNAs” network (48 nodes and 95 edges; Figure 3).

**Fig3.**
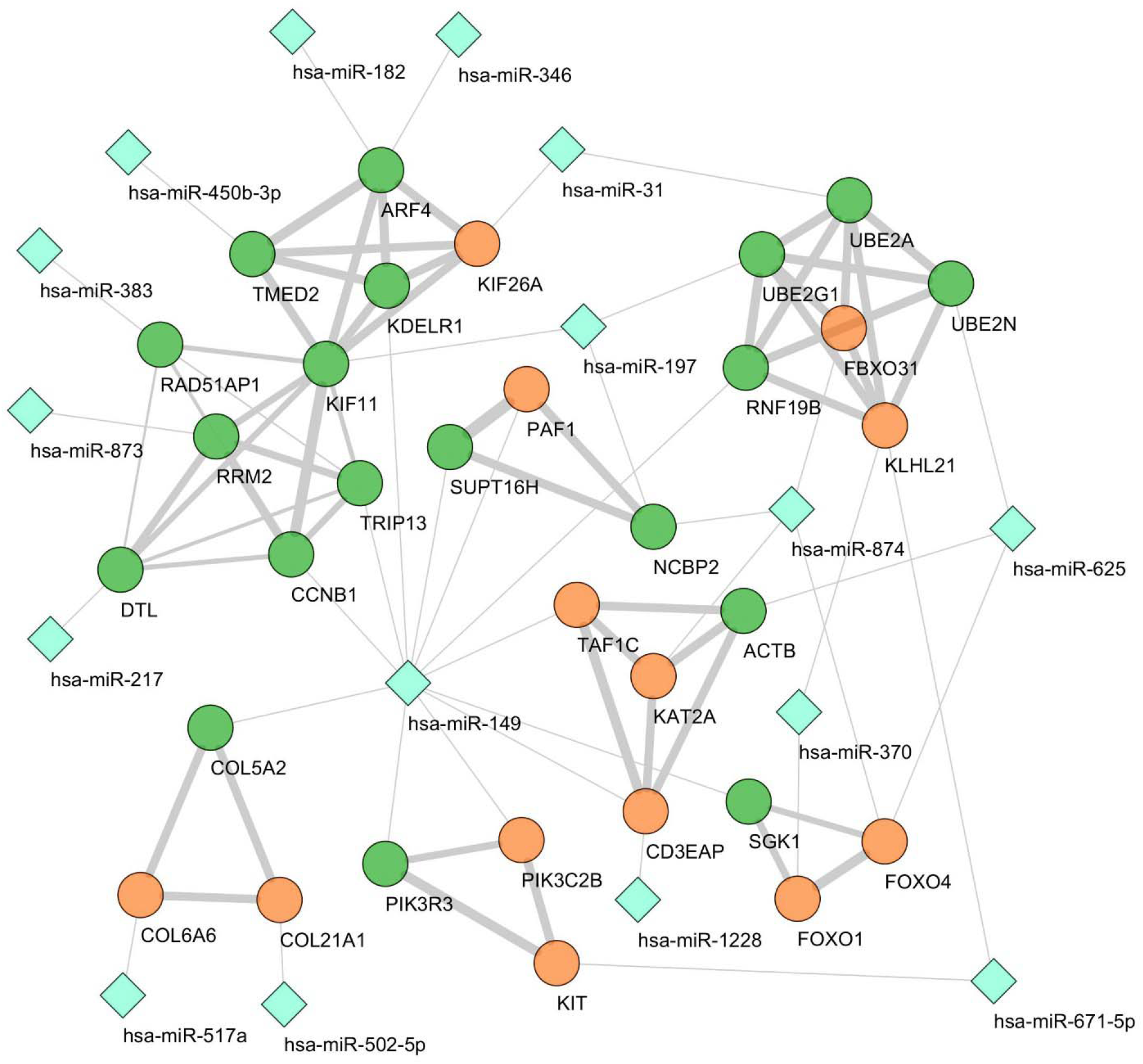
The hub clusters of PPI network of BC DEGs identified and visualized by MCODE plugin of Cytoscape 3.6.0. The corresponding miRNAs also were merged to form an informative hub mRNA-miRNA network. The blue diamond shaped, green, and orange nodes represent miRNAs, up-regulated, and down-regulated mRNAs respectively. The expression status of DEGs were obtained from microarray analysis in the current study.

Enrichment analysis including KEGG pathway and GO annotation analyses were implemented by DAVID 6.7 upon hub genes. The GO terms with *p-*value less than 0.05 counted as significantly important and the top 5 of each term were illustrated and brought in Figure 4. The most enriched pathways from KEGG was Focal adhesion (hsa04510). The most enriched BP, MF, and CC were ‘response to DNA damage stimulus; GO:0006974’, ‘ATP binding; GO:0005524’, and ‘nucleoplasm; GO:0005654’, respectively. The introduced terms were also brought in along related genes and *p-*values (Supplementary 4(B)).

**Fig4.**
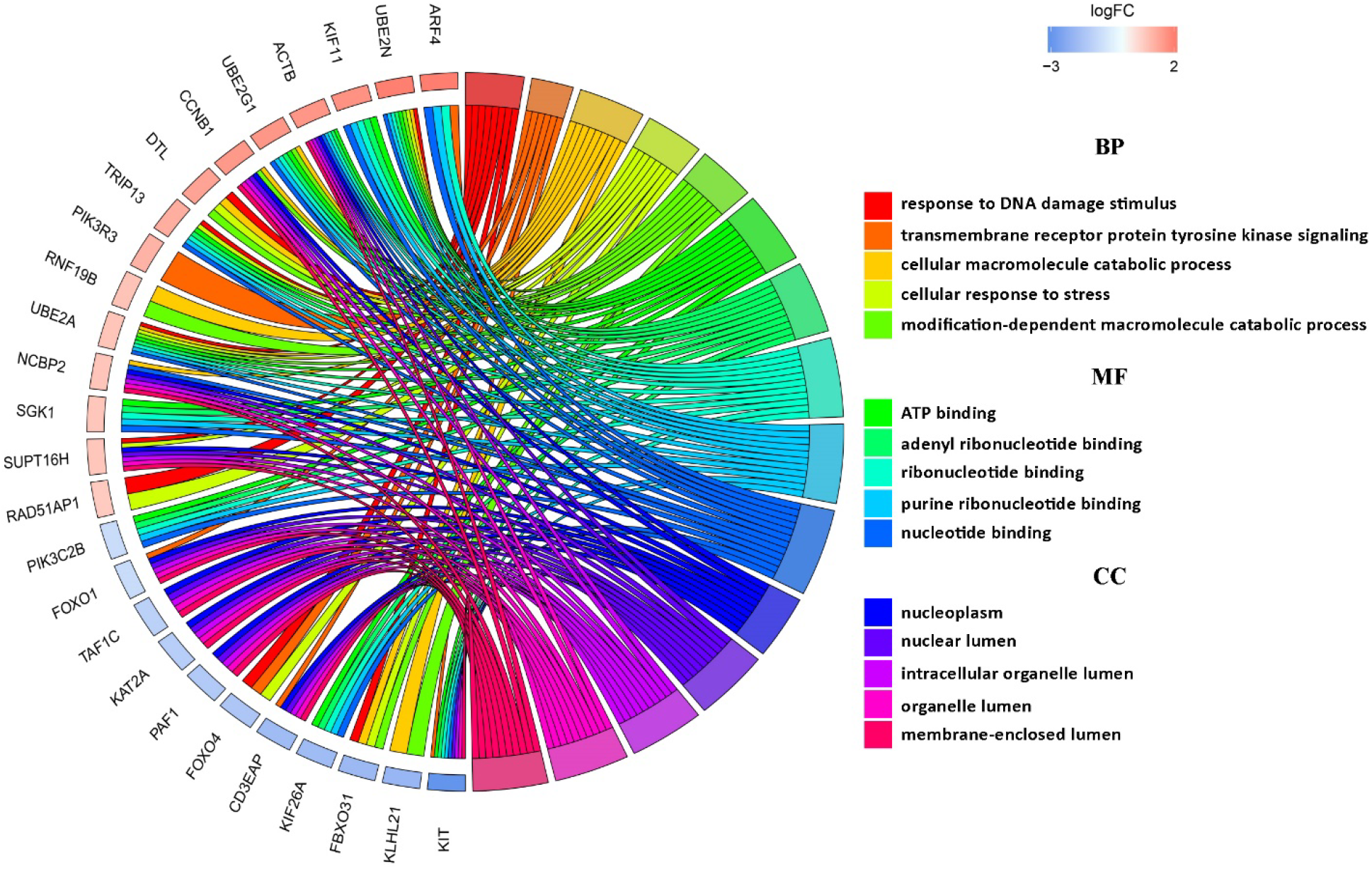
The top five enriched GO terms of hub clusters that were plotted by Goplot package of R. The BP, MF, and CC are stands for biological process, molecular function, and cell compartment respectively. The colors were explained in the legends right side of the graphs. The genes were sorted according to their logFC.

### 5. Explored MREs and Enriched Pathways and GO Terms

The networks of “hub mRNAs-miRNAs” and “hub circRNAs-hub miRNAs” were integrated together to find the common miRNAs. According to miRNA Differential Expression tool of starBase Pan-Cancer Analysis Platform, only four (hsa-miR-149, hsa-miR-182, hsa-miR-873, and hsa-miR-383) out of five common miRNAs were expressed significantly different between normal and breast cancer tissues excluding has-mir-31 (Table 2).

**Table 2.** The common hub miRNAs in BC between hub miRNA-hub circRNA and miRNA-mRNA networks that were investigated for being differentially expressed via starBase Pan-Cancer Analysis Platform RNA-seq data concerning *p-value*. The hsa-miR-31 was excluded from the list.

| Leading miRNAs | Cancer | normal | cancer Exp | nor Exp | Fold | *p-value* |
| --- | --- | --- | --- | --- | --- | --- |
| hsa-miR-149 | 1085 | 104 | 1.07 | 0.13 | 8.38 | 4.9e-20 |
| hsa-miR-182 | 1085 | 104 | 48371.55 | 9796.84 | 4.94 | 2.1e-86 |
| hsa-miR-31 | 1085 | 104 | 5.69 | 4.03 | 1.41 | 0.41 |
| hsa-miR-383 | 1085 | 104 | 1.67 | 4.30 | 0.39 | 3.4e-43 |
| hsa-miR-873 | 1085 | 104 | 0.04 | 0.01 | 4.16 | 0.017 |

The MREs were explored and revealed for leading miRNAs and hub circRNAs through Circular RNA Interactome database (Figure 5 and Supplementary 4(C)). The enriched pathways and GO terms for the leading miRNAs were also excavated and introduced (Supplementary 4(D)).

**Fig5.**
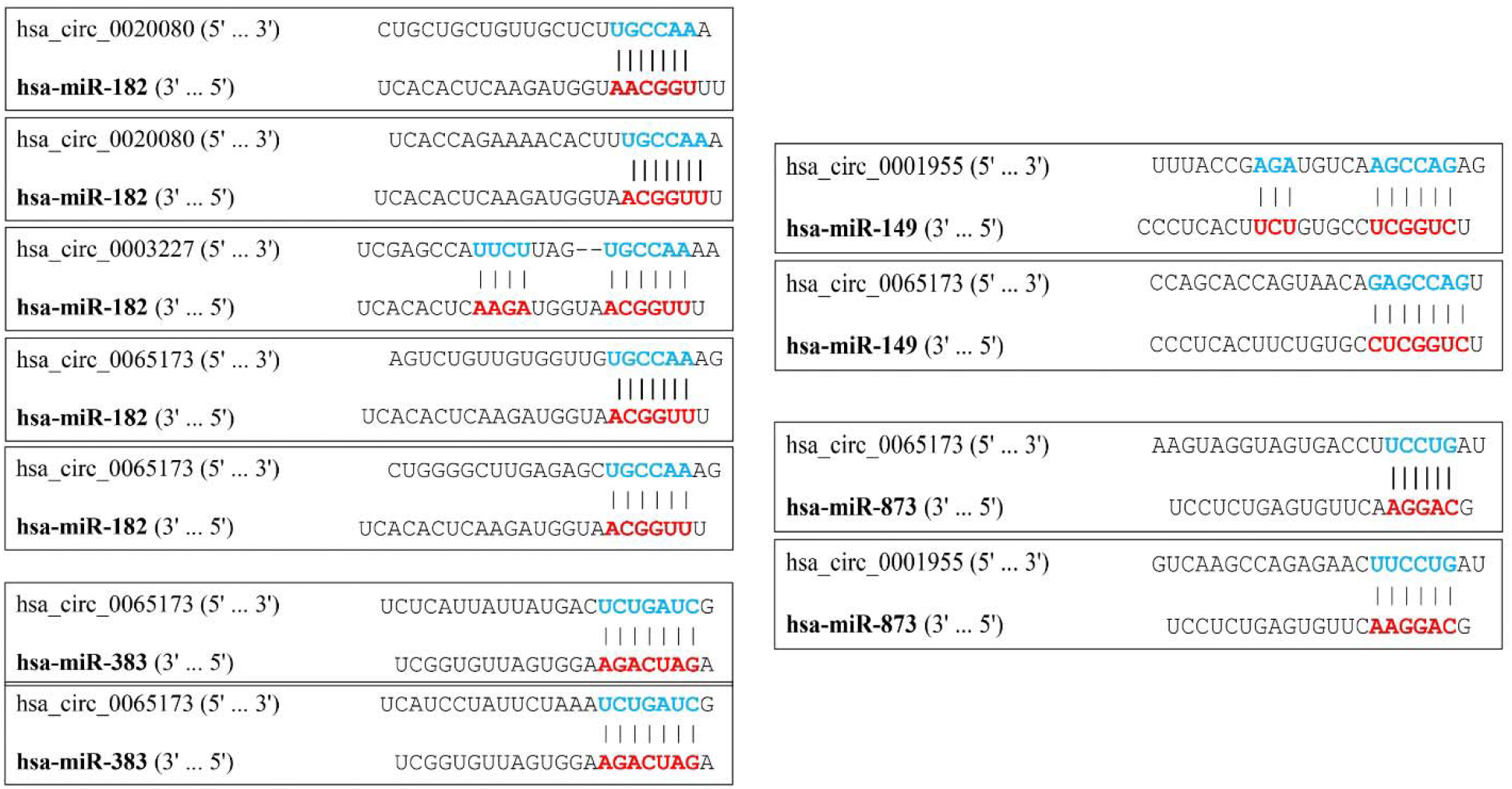
The explored MREs between hub circRNAs and leading miRNAs via Circular RNA Interactome database. The matching sequences upon hub circRNAs and leading miRNAs are respectively in blue and red colors.

## DISCUSSION

RNA splicing eventuates in circular and linear ways producing two kinds of RNAs namely, linear and circular RNAs. Linear RNAs are the result of linear splicing when introns will be separated and the exons joined together, but the back splicing culminates in circRNAs that have diverse lengths regarding have both introns and one or multiple exons [26,27]. The high throughput methods played major roles in locating and introducing novel circRNAs that count as the paramount regulators of cellular signaling pathways since they are the upstream elements controlling downstream miRNAs and mRNAs [28]. The circRNA-miRNA-mRNA axis lately engaged several oncologists, despite acting as enhancers or silencers, especially in BC for both diagnosis and treatment [29]. In the current study, we analyzed and introduced luminal A and TN common co-expressed circRNAs by using high throughput microarray data available in GEO repository. The common co-expressed elements are meticulous targets that have the chance of heterogeneity elimination which is a very delicate dilemma in BC uprooting [30]. The aforementioned subtypes also consider as the most important ones in BC, since luminal A has the title of “the most engaging” subtype and TN subtype to not have any hormonal characteristics that is hard to both cure and diagnosis [31].

The ceRNAs unite to lay the foundations of regulatory networks that affect any biological processes. The circRNAs in precise need additional investigations due to their importance regarding their abundance and ambiguity roles in breast and other human cancers. The main issue is finding the hub circRNAs that have extensive regulatory branches and in this way, avoid pathway compensation ruled by cancer cells in case of manipulating single circRNAs [32]. In the current study, the RNA-seq expression status of circRNAs corresponding genes was used for narrowing down the discovered list. Pertaining to this fact that there are some genes that are unable to produce circRNAs [33], the corresponding genes seem important both for anti-cancer treatments and diagnosis; especially because of some circRNAs functions as competitors of related mRNAs [34]. All the discovered hub circRNAs were regarded as ecRNAs, capable of regulating gene expression at both levels of post- and at transcription through sponging miRNAs [6]. According to the predicted MREs by Circular RNA Interactome, several 7- and 8-mer matches were found between leading miRNAs and hub circRNAs that promise secure and stable connection for sponging miRNAs and sequential regulation. This database takes advantage of Targetscan prediction algorithm that finds the probable complementary seeds on miRNAs and MREs of circRNAs [25]. With reference to this fact that alternative splicing undergoes alterations during breast cancer development, the potential matches between miRNAs and circRNAs ought to be as secure as possible to guarantee the true potential of circRNAs in sponging and regulating miRNAs [35,36]. The ratio of affinity to abundance plays a major role in the MREs profile explaining that despite the higher abundance of 6-mer matches in comparison to 7- and 8-mer matches, the latter ones are more vital for maintaining the connection since they generate better affinity [37,38]. In connection with this fact that ecRNAs are the only type of circRNAs that has MREs, ciRNAs and EIciRNAs serve as post-transcriptional regulators by modulating mRNA transcription and protein translation [28].

Through literature search it was found that among all the discovered RBPs, as far as this group knows, the empirical mechanisms’ investigation of EWSR1 and EIF4A3 for BC did not take place and the others were not studied fully and even served ambiguous results in case of IGFBPs [39-47]; as for, they seem to be novel and precious targets in controlling BC especially if they will be studied along related proteins in practical networks. In connection with the enriched Reactome pathways, *Estrogen-dependent gene expression*, *TP53 Regulates Metabolic Genes*, *Pre-NOTCH Transcription and Translation*, *Regulation of MECP2 expression and activity*, *Ca^2+^ pathway*, *Regulation of RUNX1 Expression and Activity*, *MAPK6/MAPK4 signaling*, *ceRNAs regulate PTEN translation*, and *Insulin-like Growth Factor-2 mRNA Binding Proteins* specifically are related to BC and although studied before, need further care, hence their manipulation with the identified RBPs or circRNAs seem to be an intelligent approach to incapacitate BC progression.

The lethality and recurrence of BC are blamed of metastasis to distant organs, and although the exact undergoing processes of metastasis are not fully understood because of complexity, the impact of epithelial-mesenchymal transition (EMT) clearly outshines and is undeniable [48]. Overexpression of methyl-CpG-binding protein (MeCP2) widely known as an oncogenic epigenetic modulator relevant to BC and acts in line with RAS, MAPK, and PI3K pathways to the extent that its down-regulation would be compensated by RAS overexpression in TN subtype [49]. Another metastasis inducer called A disintegrin and metalloprotease domain-containing protein 12 (ADAM-12) would be silenced by the regulatory axis of Z-DNA/MeCP2/NF1. In BC, lower expression of Z-DNA forming negative regulatory element (NRE) leads to its disunion from MeCP2, hence promoting EMT related metastasis following ADAM-12 up-regulation [50]. The Notch signaling cooperates diverse panels of all the cancer-promoting pathways (BC stem cell, Hedgehog, Wnt, HER/ErbB, PDGF/PDGFR, TGF-β and etc.), inflammatory and angiogenesis pathways (NF-κB signaling, Leptin signaling, EGF/VEGFR-2 signaling, and HIFκ signaling) and all cellular processes [51]. Up-regulation of Notch receptors (Notch1, Notch3, and Notch4) culminates in BC progression engaging Notch1-JAG1, RAS, and TGF-β [52-55]. In connection with the aforementioned vast characteristics of the Notch signaling one can reasonably conclude the value of its targeting by modulating every means to control BC.

The RBPs are also beneficial for targeting Runt-related transcription factor 1 (RUNX1) Expression and Activity and Ca^2+^ pathways. RUNX1 is considered to be a double-edged sword since both its up-regulation and down-regulation is detectable in BC. Loss of function mutation in cell-specific RUNX1 in luminal subtypes of BC accompanied with TGF-β leads to EMT metastasis [56]; on the other hand, its up-regulation in TN subtype triggers cancerous cells migration [57]. The Ca^2+^ pathway is also another BC triggering player (migration and angiogenesis, proliferation and apoptosis) that needs further and deeper study due to being subtype-specific. Regarding the existing gradient of Ca^2+^ on both sides of the plasma membrane and the necessity of upper Ca^2+^ concentration in the cytoplasmic environment to instigate proliferation and gene expression, the calcium exchangers and channels or the whole signaling network are advantageous to control BC [58]. It was found that targeting Secretory Pathway Ca^2+^-ATPase 1 (SPCA1) up-regulation led to proliferation subsiding as it subsequently has an impact on IGF1R production in BC basal-like subtype [59]. In the case of ER^+^ subtypes, it was approved that focusing on calcium pump plasma membrane Ca^2+^-ATPase 2 (PMCA2) overexpression culminates in breaking the apoptosis resistance [60]. These findings strengthen this idea that by modulating circRNAs and integrating the elements of converging parallel mechanisms a multi-aspects procedure would be flourished to eliminate BC.

With reference to the enriched pathways of leading miRNAs and mRNAs clusters, lots of BC related pathways were observed that emphasized the significance of discovered regulatory ceRNA axes through the current study for BC eradication. According to this disclosure, there was a common pathway among them; focal adhesion (FA). FAs are the plasma membrane inner structures that contain lots of constitutive proteins including kinases, adaptors, and phosphatases that will be congregated as the result of cells attachment to ECM. FAs are responsible for a wide variety of cellular processes from growth and differentiation to morphology, but their importance in migration is exemplary which is a result of its turn-over dominance to broadening [61]. It was demonstrated that focal adhesion kinase (FAK), which is a key player of this pathway, along being accountable for BC metastasis is also crucial for cell cycle progression engaging a myriad of elements and pathways like MAPK/ERK, Src, and cyclin D1 [62,63]. It was also determined that FAs are engrossed in generating and maintenance of breast cancer stem cells (BCSCs) since mammary gland stem cells (MaSCs) have a panel of overexpressed integrins that follows by FAK activation, anoikis evading (programmed cell death that is the result of detaching from ECM [64]), and self-renewal emergence [65,66].

## CONCLUSION

This paper has investigated the high-priority role of ceRNAs regulatory networks in BC and led us to conclude that manipulating the circRNA-miRNA-mRNA or RBP-circRNA-miRNA axes potentially culminates in halting several BC triggering biological processes and pathways. We have found several novel controllable hubs and targets including circRNAs, miRNAs, RBPs, and mRNAs. The introduced cutting-edge strategy, which is targeting upper hand elements instead of just controlling downstream mRNAs and proteins, needs further wet lab experiments that this group is now working on it. Furthermore, the introduced regulatory elements that were obtained through merging several promising bioinformatics tools serve as individual valuable particles to devise inhibitory procedures against them. The study also covered the explanation of some salient and fundamental pathways in BC and explained the probability effect of controlling RBPs and circRNAs on them, hence suggesting another aspect of ceRNAs networks’ regulation.

## AUTHORS CONTRIBUTIONS

M.S conceptualized and supervised the project. F.A performed and analyzed all the data and wrote the manuscript. M.S and F.A have deciphered the analyzed data, read, and edited the manuscript.

## CONFLICTS OF INTEREST

The author(s) declared no potential conflicts of interest with respect to the research, authorship, and/or publication of this article.

## FUNDING

This research did not receive any grant from funding agencies in the public, commercial, or not-for-profit sectors.

## REFERENCES

1. Siegel RL, Miller KD, Jemal A (2017) Cancer statistics, 2017. CA: a cancer journal for clinicians 67 (1):7–30

2. Salmena L, Poliseno L, Tay Y, Kats L, Pandolfi PP (2011) A ceRNA hypothesis: the Rosetta Stone of a hidden RNA language? Cell 146 (3):353–358

3. Bosson AD, Zamudio JR, Sharp PA (2014) Endogenous miRNA and target concentrations determine susceptibility to potential ceRNA competition. Molecular cell 56 (3):347–359

4. Yang Y, Fan X, Mao M, Song X, Wu P, Zhang Y, Jin Y, Yang Y, Chen L-L, Wang Y (2017) Extensive translation of circular RNAs driven by N 6-methyladenosine. Cell research 27 (5):626

5. Danan M, Schwartz S, Edelheit S, Sorek R (2011) Transcriptome-wide discovery of circular RNAs in Archaea. Nucleic acids research 40 (7):3131–3142

6. Zhang X-O, Wang H-B, Zhang Y, Lu X, Chen L-L, Yang L (2014) Complementary sequence-mediated exon circularization. Cell 159 (1):134–147

7. Zhang Y, Zhang X-O, Chen T, Xiang J-F, Yin Q-F, Xing Y-H, Zhu S, Yang L, Chen L-L (2013) Circular intronic long noncoding RNAs. Molecular cell 51 (6):792–806

8. Li Z, Huang C, Bao C, Chen L, Lin M, Wang X, Zhong G, Yu B, Hu W, Dai L (2015) Exon-intron circular RNAs regulate transcription in the nucleus. Nature structural & molecular biology 22 (3):256

9. Xu S, Zhou L, Ponnusamy M, Zhang L, Dong Y, Zhang Y, Wang Q, Liu J, Wang K (2018) A comprehensive review of circRNA: From purification and identification to disease marker potential. PeerJ 6:e5503

10. Lü L, Sun J, Shi P, Kong W, Xu K, He B, Zhang S, Wang J (2017) Identification of circular RNAs as a promising new class of diagnostic biomarkers for human breast cancer. Oncotarget 8 (27):44096

11. Wu J, Jiang Z, Chen C, Hu Q, Fu Z, Chen J, Wang Z, Wang Q, Li A, Marks JR (2018) CircIRAK3 sponges miR-3607 to facilitate breast cancer metastasis. Cancer letters

12. Liang H-F, Zhang X-Z, Liu B-G, Jia G-T, Li W-L (2017) Circular RNA circ-ABCB10 promotes breast cancer proliferation and progression through sponging miR-1271. American journal of cancer research 7 (7):1566

13. Bartel DP (2009) MicroRNAs: target recognition and regulatory functions. cell 136 (2):215–233

14. Cai Y, Yu X, Hu S, Yu J (2009) A brief review on the mechanisms of miRNA regulation. Genomics, proteomics & bioinformatics 7 (4):147–154

15. O’Day E, Lal A (2010) MicroRNAs and their target gene networks in breast cancer. Breast cancer research 12 (2):201

16. Imani S, Wei C, Cheng J, Khan MA, Fu S, Yang L, Tania M, Zhang X, Xiao X, Zhang X (2017) MicroRNA-34a targets epithelial to mesenchymal transition-inducing transcription factors (EMT-TFs) and inhibits breast cancer cell migration and invasion. Oncotarget 8 (13):21362

17. Li P, Sheng C, Huang L, Zhang H, Huang L, Cheng Z, Zhu Q (2014) MiR-183/-96/-182 cluster is up-regulated in most breast cancers and increases cell proliferation and migration. Breast cancer research 16 (6):473

18. Abdelmohsen K, Gorospe M (2010) Posttranscriptional regulation of cancer traits by HuR. Wiley Interdisciplinary Reviews: RNA 1 (2):214–229

19. Hentze MW, Preiss T (2013) Circular RNAs: splicing’s enigma variations. The EMBO journal 32 (7):923–925

20. Lécuyer E, Yoshida H, Parthasarathy N, Alm C, Babak T, Cerovina T, Hughes TR, Tomancak P, Krause HM (2007) Global analysis of mRNA localization reveals a prominent role in organizing cellular architecture and function. Cell 131 (1):174–187

21. Glisovic T, Bachorik JL, Yong J, Dreyfuss G (2008) RNA-binding proteins and post-transcriptional gene regulation. FEBS letters 582 (14):1977–1986

22. Zheng L, Zhang Z, Zhang S, Guo Q, Zhang F, Gao L, Ni H, Guo X, Xiang C, Xi T (2018) RNA Binding Protein RNPC1 Inhibits Breast Cancer Cell Metastasis via Activating STARD13-Correlated ceRNA Network. Molecular pharmaceutics 15 (6):2123–2132

23. Guo X, Hartley RS (2006) HuR contributes to cyclin E1 deregulation in MCF-7 breast cancer cells. Cancer research 66 (16):7948–7956

24. Maubant S, Tesson B, Maire V, Ye M, Rigaill G, Gentien D, Cruzalegui F, Tucker GC, Roman-Roman S, Dubois T (2015) Transcriptome analysis of Wnt3a-treated triple-negative breast cancer cells. PloS one 10 (4):e0122333

25. Dudekula DB, Panda AC, Grammatikakis I, De S, Abdelmohsen K, Gorospe M (2016) CircInteractome: a web tool for exploring circular RNAs and their interacting proteins and microRNAs. RNA biology 13 (1):34–42

26. Gao Y, Zhao F (2018) Computational strategies for exploring circular RNAs. Trends in Genetics 34 (5):389–400

27. Li X, Chu C, Pei J, Măndoiu I, Wu Y (2018) CircMarker: a fast and accurate algorithm for circular RNA detection. BMC genomics 19 (6):175

28. Rong D, Sun H, Li Z, Liu S, Dong C, Fu K, Tang W, Cao H (2017) An emerging function of circRNA-miRNAs-mRNA axis in human diseases. Oncotarget 8 (42):73271

29. Nair AA, Niu N, Tang X, Thompson KJ, Wang L, Kocher J-P, Subramanian S, Kalari KR (2016) Circular RNAs and their associations with breast cancer subtypes. Oncotarget 7 (49):80967

30. Schiavon G, Smid M, Gupta GP, Redana S, Santini D, Martens JW (2012) Heterogeneity of breast cancer: gene signatures and beyond. In: Diagnostic, Prognostic and Therapeutic Value of Gene Signatures. Springer, pp 13–25

31. Kennecke H, Yerushalmi R, Woods R, Cheang MCU, Voduc D, Speers CH, Nielsen TO, Gelmon K (2010) Metastatic behavior of breast cancer subtypes. Journal of clinical oncology 28 (20):3271–3277

32. Giza DE, Vasilescu C, Calin GA (2014) MicroRNAs and ceRNAs: therapeutic implications of RNA networks. Expert opinion on biological therapy 14 (9):1285–1293

33. Ivanov A, Memczak S, Wyler E, Torti F, Porath HT, Orejuela MR, Piechotta M, Levanon EY, Landthaler M, Dieterich C (2015) Analysis of intron sequences reveals hallmarks of circular RNA biogenesis in animals. Cell reports 10 (2):170–177

34. Ashwal-Fluss R, Meyer M, Pamudurti NR, Ivanov A, Bartok O, Hanan M, Evantal N, Memczak S, Rajewsky N, Kadener S (2014) circRNA biogenesis competes with pre-mRNA splicing. Molecular cell 56 (1):55–66

35. Venables JP, Klinck R, Koh C, Gervais-Bird J, Bramard A, Inkel L, Durand M, Couture S, Froehlich U, Lapointe E (2009) Cancer-associated regulation of alternative splicing. Nature structural & molecular biology 16 (6):670

36. Mayr C, Bartel DP (2009) Widespread shortening of 31 UTRs by alternative cleavage and polyadenylation activates oncogenes in cancer cells. Cell 138 (4):673–684

37. Grimson A, Farh KK-H, Johnston WK, Garrett-Engele P, Lim LP, Bartel DP (2007) MicroRNA targeting specificity in mammals: determinants beyond seed pairing. Molecular cell 27 (1):91–105

38. Qi X, Zhang D-H, Wu N, Xiao J-H, Wang X, Ma W (2015) ceRNA in cancer: possible functions and clinical implications. Journal of Medical Genetics 52 (10):710–718

39. Zhang Z, Huang A, Zhang A, Zhou C (2017) HuR promotes breast cancer cell proliferation and survival via binding to CDK3 mRNA. Biomedicine & Pharmacotherapy 91:788–795

40. Ke H, Zhao L, Feng X, Xu H, Zou L, Yang Q, Su X, Peng L, Jiao B (2016) NEAT1 is required for survival of breast cancer cells through FUS and miR-548. Gene regulation and systems biology 10:GRSB. S29414

41. He X, Arslan A, Ho T, Yuan C, Stampfer M, Beck W (2014) Involvement of polypyrimidine tract-binding protein (PTBP1) in maintaining breast cancer cell growth and malignant properties. Oncogenesis 3 (1):e84

42. Lucá R, Averna M, Zalfa F, Vecchi M, Bianchi F, La Fata G, Del Nonno F, Nardacci R, Bianchi M, Nuciforo P (2013) The fragile X protein binds mRNAs involved in cancer progression and modulates metastasis formation. EMBO molecular medicine 5 (10):1523–1536

43. Janowski BA, Huffman KE, Schwartz JC, Ram R, Nordsell R, Shames DS, Minna JD, Corey DR (2006) Involvement of AGO1 and AGO2 in mammalian transcriptional silencing. Nature Structural and Molecular Biology 13 (9):787

44. Patel AV, Cheng I, Canzian F, Le Marchand L, Thun MJ, Berg CD, Buring J, Calle EE, Chanock S, Clavel-Chapelon F (2008) IGF-1, IGFBP-1, and IGFBP-3 polymorphisms predict circulating IGF levels but not breast cancer risk: findings from the Breast and Prostate Cancer Cohort Consortium (BPC3). PloS one 3 (7):e2578

45. McGuire Jr WL, Jackson JG, Figueroa JA, Shimasaki S, Powell DR, Yee D (1992) Regulation of Insulin-Like Growth Factor-Binding Protein (IGFBP) Expression by Breast Cancer Cells: Use of IGFBP-1 as as Inhibitor of Insulin-like Growth Factor Action. JNCI: Journal of the National Cancer Institute 84 (17):1336–1341

46. Flint DJ, Tonner E, Allan GJ (2000) Insulin-like growth factor binding proteins: IGF-dependent and-independent effects in the mammary gland. Journal of mammary gland biology and neoplasia 5 (1):65–73

47. Yamanaka Y, Fowlkes JL, Wilson EM, Rosenfeld RG, Oh Y (1999) Characterization of insulin-like growth factor binding protein-3 (IGFBP-3) binding to human breast cancer cells: kinetics of IGFBP-3 binding and identification of receptor binding domain on the IGFBP-3 molecule. Endocrinology 140 (3):1319–1328

48. Scully OJ, Bay B-H, Yip G, Yu Y (2012) Breast cancer metastasis. Cancer Genomics-Proteomics 9 (5):311–320

49. Neupane M, Clark AP, Landini S, Birkbak NJ, Eklund AC, Lim E, Culhane AC, Barry WT, Schumacher SE, Beroukhim R (2016) MECP2 is a frequently amplified oncogene with a novel epigenetic mechanism that mimics the role of activated RAS in malignancy. Cancer discovery 6 (1):45–58

50. Ray BK, Dhar S, Henry CJ, Rich A, Ray A (2012) Epigenetic regulation by Z-DNA silencer function controls cancer-associated ADAM-12 expression in breast cancer: cross talk between MECP2 and NFI transcription factor family. Cancer research:canres. 2601.2012

51. Guo S, Liu M, Gonzalez-Perez RR (2011) Role of Notch and its oncogenic signaling crosstalk in breast cancer. Biochimica et Biophysica Acta (BBA)-Reviews on Cancer 1815 (2):197–213

52. Imatani A, Callahan R (2000) Identification of a novel NOTCH-4/INT-3 RNA species encoding an activated gene product in certain human tumor cell lines. Oncogene 19 (2):223

53. Zhang Z, Wang H, Ikeda S, Fahey F, Bielenberg D, Smits P, Hauschka PV (2010) Notch3 in human breast cancer cell lines regulates osteoblast-cancer cell interactions and osteolytic bone metastasis. The American journal of pathology 177 (3):1459–1469

54. Soares R, Balogh G, Guo S, Gartner F, Russo J, Schmitt F (2004) Evidence for the notch signaling pathway on the role of estrogen in angiogenesis. Molecular endocrinology 18 (9):2333–2343

55. Weijzen S, Rizzo P, Braid M, Vaishnav R, Jonkheer SM, Zlobin A, Osborne BA, Gottipati S, Aster JC, Hahn WC (2002) Activation of Notch-1 signaling maintains the neoplastic phenotype in human Ras-transformed cells. Nature medicine 8 (9):979

56. Hong D, Messier TL, Tye CE, Dobson JR, Fritz AJ, Sikora KR, Browne G, Stein JL, Lian JB, Stein GS (2017) Runx1 stabilizes the mammary epithelial cell phenotype and prevents epithelial to mesenchymal transition. Oncotarget 8 (11):17610

57. Browne G, Taipaleenmäki H, Bishop NM, Madasu SC, Shaw LM, Van Wijnen AJ, Stein JL, Stein GS, Lian JB (2015) Runx1 is associated with breast cancer progression in MMTV-PyMT transgenic mice and its depletion in vitro inhibits migration and invasion. Journal of cellular physiology 230 (10):2522–2532

58. Azimi I, Roberts-Thomson S, Monteith G (2014) Calcium influx pathways in breast cancer: opportunities for pharmacological intervention. British journal of pharmacology 171 (4):945–960

59. Grice DM, Vetter I, Faddy HM, Kenny PA, Roberts-Thomson SJ, Monteith GR (2010) The Golgi calcium pump secretory pathway calcium ATPase 1 (SPCA1) is a key regulator of insulin-like growth factor receptor (IGF1R) processing in the basal-like breast cancer cell line MDA-MB-231. Journal of Biological Chemistry:jbc. M110. 163329

60. VanHouten J, Sullivan C, Bazinet C, Ryoo T, Camp R, Rimm DL, Chung G, Wysolmerski J (2010) PMCA2 regulates apoptosis during mammary gland involution and predicts outcome in breast cancer. Proceedings of the National Academy of Sciences 107 (25):11405–11410

61. Nagano M, Hoshino D, Koshikawa N, Akizawa T, Seiki M (2012) Turnover of focal adhesions and cancer cell migration. International journal of cell biology 2012

62. Ghiso JAA (2002) Inhibition of FAK signaling activated by urokinase receptor induces dormancy in human carcinoma cells in vivo. Oncogene 21 (16):2513

63. Luo M, Fan H, Nagy T, Wei H, Wang C, Liu S, Wicha MS, Guan J-L (2009) Mammary epithelial-specific ablation of the focal adhesion kinase suppresses mammary tumorigenesis by affecting mammary cancer stem/progenitor cells. Cancer research 69 (2):466–474

64. Frisch SM, Screaton RA (2001) Anoikis mechanisms. Current opinion in cell biology 13 (5):555–562

65. Luo M, Guan J-L (2010) Focal adhesion kinase: a prominent determinant in breast cancer initiation, progression and metastasis. Cancer letters 289 (2):127–139

66. Frisch SM, Vuori K, Ruoslahti E, Chan-Hui P-Y (1996) Control of adhesion-dependent cell survival by focal adhesion kinase. The Journal of cell biology 134 (3):793–799

